# Enhancing SNR and generating contrast for cryo-EM images with convolutional neural networks

**DOI:** 10.1101/2020.08.16.253070

**Authors:** Eugene Palovcak, Daniel Asarnow, Melody G. Campbell, Zanlin Yu, Yifan Cheng

**Affiliations:** Department of Biochemistry and Biophysics, University of California San Francisco, San Francisco, CA 94143; Howard Hughes Medical Institute, University of California San Francisco, San Francisco, CA 94132

**Keywords:** cryo-EM, deep learning, denoise

## Abstract

In cryogenic electron microscopy (cryo-EM) of radiation-sensitive biological samples, both the signal-to-noise ratio (SNR) and the contrast of images are critically important in the image processing pipeline. Classic methods improve low-frequency image contrast experimentally, by imaging with high defocus, or computationally, by applying various types of low-pass filter. These contrast improvements typically come at the expense of high-frequency SNR, which is suppressed by high-defocus imaging and removed by low pass filtration. Here, we demonstrate that a convolutional neural network (CNN) denoising algorithm can be used to significantly enhance SNR and generate contrast in cryo-EM images. We provide a quantitative evaluation of bias introduced by the denoising procedure and its influences on image processing and three-dimensional reconstructions. Our study suggests that besides enhancing the visual contrast of cryo-EM images, the enhanced SNR of denoised images may facilitate better outcomes in the other parts of the image processing pipeline, such as classification and 3D alignment. Overall, our results provide a ground of using denoising CNNs in the cryo-EM image processing pipeline.

## Introduction

In single particle cryogenic electron microscopy (cryo-EM), high-resolution three-dimensional (3D) structures of biological macromolecules are determined by iteratively aligning and averaging a large number of very noisy two-dimensional (2D) projection images of molecules embedded in a thin layer of vitreous ice (Cheng, 2015). This process requires identification of individual particles in brightfield images (particle picking), sorting the particle images according to conformational state (classification), iteratively determining the orientation of each particle (alignment) and, finally, calculating a 3D reconstruction. The success of each of these steps is critically dependent on the signal-to-noise ratio (SNR) of the cryo-EM images at all frequencies (Jensen, 2001). The frequency dependent SNR is mathematically well defined (Bershad and Rockmore, 1974; Frank and Alali, 1975). The visual contrast of cryo-EM images is less strictly defined but is closely related to the low frequency amplitudes. Both are fundamentally limited by the radiation sensitivity and transparency of the frozen-hydrated cryo-EM specimen, which necessitates low-dose phase-contrast imaging (Glaeser, 1999). They are further limited by the shape of the contrast transfer function (CTF), which is a sine function and suppresses the amplitude at the low-spatial frequencies responsible for producing image contrast (Wade, 1992).

Within these constraints, the SNR of cryo-EM images is typically well below 1 (more noise than signal) (Frank and Alali, 1975). The conventional approach to maximizing low-frequency SNR and increasing image contrast is to increase the defocus of the objective lens. Higher defocus alters the CTF of the electron microscope and produces higher amplitudes thus more contrast and SNR at low spatial frequencies, but at the expenses of reducing amplitude (thus SNR) at the intermediate and high-spatial frequencies required for high-resolution structure determination (Cheng, 2015). This intrinsic issue becomes more acute for small or irregularly shaped particles and greatly increases the difficulty of determining the 3D structures of such specimens (Herzik et al., 2019). Another method to generate image contrast is to use a phase plate to generate a phase shift in CTF, thus to improve the contrast at low spatial frequencies without perturbing the information at high spatial frequencies (Danev and Nagayama, 2001). The most successful phase plate device currently available is the Volta phase plate, which is a continuous carbon film placed in the back-focal plane (Danev et al., 2017). In practice, however, the Volta phase plate causes a noticeable loss of SNR at high-frequency, likely due to the “absorption” of electrons by the carbon film (personal communication with Rado Danev). A new type of laser phase plate being currently developed will presumably not have such issues (Schwartz et al., 2019; Schwartz et al., 2018). Nevertheless, complementary and alternative methods of image contrast enhancement in cryo-EM could therefore be of great value.

Here, we present a computational image restoration approach to reduce the noise that limits SNR and contrast in cryo-EM images. The basic idea is to train a parameterized image operator (a convolutional neural network or CNN) as an image denoiser. The training scheme used in our approach, called *noise2noise* (Lehtinen et al., 2018), uses noisy cryo-EM data as a training signal and is fully compatible with existing strategies for cryo-EM data acquisition. We show that CNNs trained with *noise2noise* significantly enhance the contrast of cryo-EM images, similar to the effects of the Volta phase plate. In terms of SNR, we found that the denoising CNNs greatly reduce the noise power at all spatial frequencies. At low and intermediate spatial frequencies, this corresponds to a genuine increase in the relative strength of the true signal in the image. At higher spatial frequencies, noise reduction by CNNs also introduces false signal or ‘bias’. We developed methods to quantify such bias induced by the denoiser and evaluated with examples the influences such bias on image alignment and 3D reconstruction. We show that the bias introduced by CNNs during the denoising process does not prevent highly accurate alignment of denoised particles and largely averages out during 3D reconstructions. Likely, practical and broad use of denoised images in all stages of the cryo-EM image processing pipeline will require some minor adaptations of existing 3D reconstruction software. However, our results demonstrate that there is no reason, in principle, that denoising CNNs could not be used to great benefit in single-particle cryo-EM.

While we were preparing this work, several other groups have introduced *noise2noise*-based denoising CNNs for enhancing contrast of cryo-EM images (Bepler et al., 2019a; Bepler et al., 2019b; Tegunov and Cramer, 2019). The application so far is, however, limited to the level of visualization and particle picking. Our quantitative characterizations of the influence of denoising CNNs on the SNR broaden the potential applications of denoising CNNs in single particle cryo-EM image processing pipeline.

## Results

### Training a convolutional neural network to denoise cryo-EM images

CNNs are powerful parameterized function approximators (Krizhevsky et al., 2017). They consist of a large number of small convolution filters with learnable parameters. Each convolutional filter is applied to the input image in real-space and passed through a pixel-wise nonlinear function with a shape parameter (typically a threshold-based masking function called a ‘rectified linear unit’ or ‘ReLU’). By stacking a large number of these simple operations in series (convolutional layers), complex input-output maps (functions) can be approximated (Goodfellow et al., 2016). Since each operation in a CNN is differentiable, the parameters of a CNN can be learned from a large set of input-output image pairs using gradient-based stochastic optimization. To train a CNN to approximate an image denoiser, these input-output pairs typically consist of an image with and without noise (Zhang et al., 2017). Training consists of calibrating the CNN’s parameters such that applying the CNN to the noisy image produces the noiseless image. To ensure that the CNN generalizes properly to unseen images, the content of the images and the distribution of the noise in the training dataset must be representative of those expected during use. These requirements, however, pose a major problem for training a denoising CNN for cryo-EM images: the radiation sensitivity of frozen-hydrated biological samples makes it impossible to obtain a noiseless cryo-EM image *in principle* (Glaeser, 1999).

Lehtinen et al. recently demonstrated that pairs of noisy images of the same object can be used in place of noisy/noiseless pairs to train denoising CNNs (Lehtinen et al., 2018). Training is performed as if the noisy image pair were a noisy/noiseless pair: the CNN is applied to the first noisy image and the discrepancy between the output and the second noisy image is calculated (the loss). The loss is used to calculate the direction each parameter should be optimized to make the output and second noisy image more similar (the loss gradient). Lehtinen et al. showed that given a sufficiently large set of noisy image pairs, the parameters of the denoiser converge on the same parameter values that would be obtained using conventional noisy/noiseless training data.

This training strategy, termed *noise2noise*, enables the parameters of a denoising CNN to be learned without ever seeing noiseless images. It simply requires that the noise in the training pairs be statistically uncorrelated given the signal. In fact, *noise2noise* training makes no other assumptions about the structure of the signal or the distribution of the noise and learns both implicitly from the training data. A more mathematical description of the *noise2noise* training scheme is provided in the supplement and we also refer the reader to the original *noise2noise* paper and the more general framework for self-supervised denoising described in *noise2self* (Supplemental note 1, and (Batson and Royer, 2019; Lehtinen et al., 2018)).

Nowadays, cryo-EM images recorded using direct electron detection cameras are dose-fractionated movies containing many frames. We can generate two images with the same signal but uncorrelated noise by summing up disjoint sets of movie frames, such as the even and odd frames after motion correction (Zheng et al., 2017) and electron dose-weighting (Grant and Grigorieff, 2015). Applying the *noise2noise* scheme to cryo-EM images therefore requires no alteration in data collection and can work on previously collected datasets.

Note that the noise considered here is mostly shot noise and not amorphous features in the sample such as particle debris or the background added by vitreous ice. Such features are considered ‘structural noise’ when calculating 3D reconstructions, but the denoising CNN has no way of distinguishing them from the desired particle signal and cannot remove them. Nevertheless, shot noise is the dominant source of noise in cryo-EM imaging (Baxter et al., 2009).

### Implementing a denoising CNN for cryo-EM

In principle, the *noise2noise* training scheme only requires a large dataset of noisy image pairs to train a denoising CNN. In practice, we found that denoising performance depends strongly on the structure of the CNN and the preprocessing of the training images. We implemented *noise2noise* procedure in a denoising program, *restore*, in which we used a CNN architecture similar to the U-net used by Lehtinen et al. but with some modifications (Ronneberger et al., 2015). We replaced each block of convolutional layers in the U-net with a wide-activation convolutional layer (Yu et al., 2018), and used depth-to-space up-sampling to minimize aliasing artifacts (Odena et al., 2016). We found these modifications improved the training loss and the visual quality of the output compared to a standard U-net. In the supplement, we provide a full description of the CNN architecture used in this work (Supplemental figure 2, Supplemental note 1).

Currently, for each specific cryo-EM dataset, we use part of the dataset as training pairs to train a specific CNN and use the trained CNN to denoise cryo-EM images of the entire dataset. This is possible because a typical cryo-EM dataset consists of thousands of movies and the training process is relatively fast (a few hours). With such an abundance of training data, overfitting the CNN is very unlikely.

To generate training data from a set of dose-fractionated electron movies, we first generate motion-corrected and dose-weighted sums of the even and odd frames using MotionCor2 (Zheng et al., 2017). We have tested our procedure in a number of examples (discussed below). The physical pixel sizes of these datasets are approximately 1Å. Each cryo-EM image is modified by the CTF of the electron microscope. Considering that the defocus and astigmatism of each image is different, the CTF modifies otherwise similar signals in unique but predictable ways and a denoising CNN would need to learn to distinguish all CTF-aberrated signals from noise. We reasoned that it would be advantageous to correct the CTF in the noisy images upfront, so that the CNN would not need to learn to be invariant to CTF modulations of the underlying signal. Thus, before training and applying the CNN to denoise all images in a dataset, we performed phase-flipping on the images by multiplying their Fourier transforms with the sign of the CTF and computing the inverse Fourier transform. While the examples we show in the following are phase-flipped prior training CNN and denoising image, it may be unnecessary to phase-flip the training images prior to the denoising procedure.

Considering that SNR at very high spatial frequencies is very low (Baxter et al., 2009), we do not expect any denoising algorithm to recover the signal under such conditions. We typically bin training images by Fourier cropping, which effectively reduces the total noise in the image and makes it smaller. In the example of the 20S proteasome shown later, we Fourier crop the training images to a pixel size ~1.5Å. To facilitate training in batches, we break up the phase-flipped, Fourier-cropped training images into square patches (typically 192×192 pixels). For most macromolecular specimens examined, a patch of this size contains at least several particles. We normalize each patch by subtracting the mean pixel value and dividing by the standard deviation.

Finally, we train the CNN using the Adam stochastic optimization algorithm with weight normalization (Kingma and Ba, 2014; Salimans and Kingma, 2016). We find that training CNNs with *noise2noise* is fast, typically achieving a stable plateau in the value of the loss function in several epochs (20-30 minutes on a single modern GPU). We train for one hundred epochs.

Once trained, the CNN can be used to denoise full images. We preprocess these images in the same way we did during training, except we do not break the image into patches. This is possible because CNNs can operate on images with arbitrary shape and size. Finally, we unbin the denoised image by zero-padding its Fourier transform, low-pass filtering with a soft radial Fourier mask, and calculating the inverse Fourier transform. The result is a denoised image with the shape and pixel size of the original image, bandlimited to the same frequency as the binned training data.

### Contrast enhancement for diverse cryo-EM specimens

We trained denoising CNNs on several previously collected cryo-EM datasets with a range of specimen types and imaging conditions: the Falciparum 80S ribosome (~3.2 MDa), the human TRPM4 ion channel (~700 kDa), a human integrin-Fab complex (~280 kDa), and the human PKA catalytic domain (~43 kDa). Despite differences in particle shape, molecular weight, and imaging conditions, all denoised images show significantly enhanced visual contrast with particle shapes are clearly defined and consistent with the shape of these molecules (Figure 1A-D) (Autzen et al., 2018; Campbell et al., 2020; Herzik et al., 2019; Wong et al., 2014). We show four cases here, but we have not found any case where the contrast of single-particle cryo-EM specimens is not significantly enhanced by training and applying a denoising CNN.

**Figure 1.**
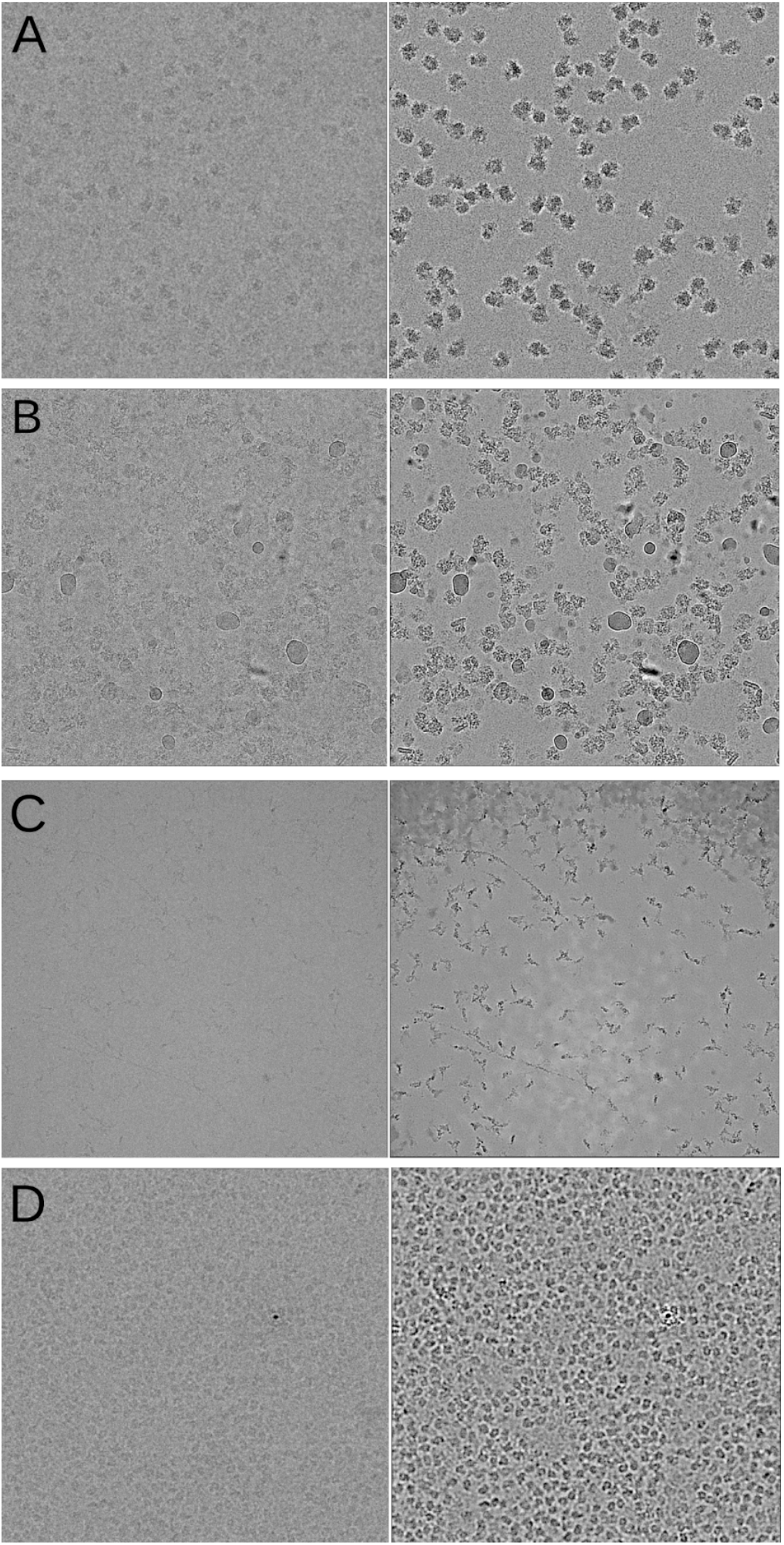
Performance of denoising CNNs on real cryo-EM images. (**A**) Falciparum ribosome particles, (**B**) TRPM4 particles, (**C**) integrin-Fab particles, (**D**) protein kinase A particles, before (left) and after (right) denoising.

To understand how a denoising CNN affects information at different spatial frequencies in the image, we trained a denoising CNN on a dataset of a well-characterized test specimen, the *Thermoplasma acidophilum* 20S proteasome (T20S) (Figure 2). For this dataset, we used all images in the dataset to train the denoiser and we bandlimited the resolution of training data to 3Å by Fourier cropping. Compared with the original image (Figure 2A), denoised image (Figure 2B) shows significant image contrast without blurring shown in low pass filtered image (Figure 2C).

**Figure 2.**
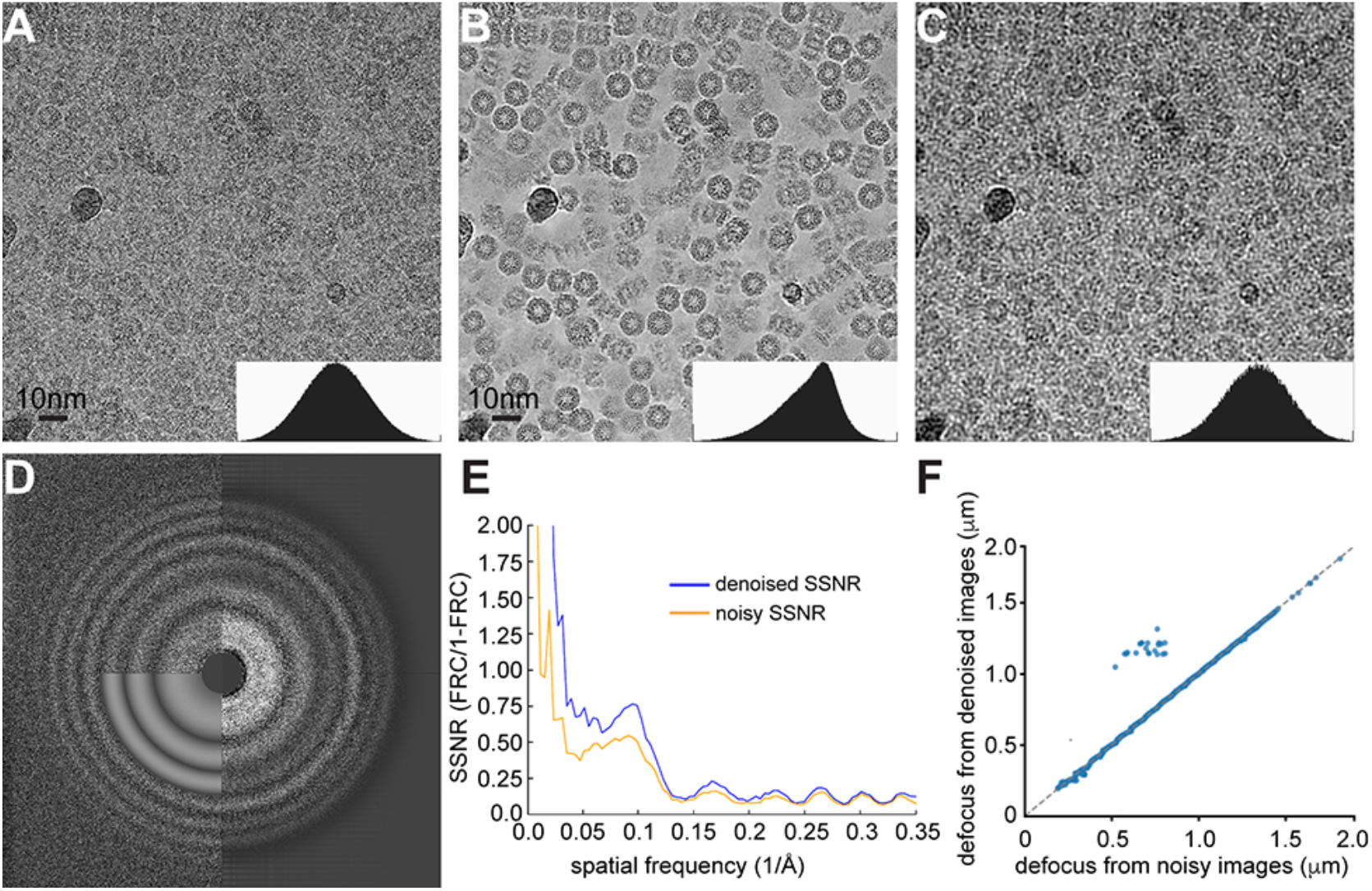
Effects of denoising on Fourier amplitudes. (**A**) Cryo-EM image of 20S proteasome recorded at a defocus of −0.4μm. (**B**) Same cryo-EM image after denoising. (**C**) Same image after applying a low pass filtered in Fourier space to 1/20Å. Image contrast in A, B and C are manually scaled so that the histograms of pixel intensities are similar (small insert in each panel). (**D**) Fourier transforms calculated from original (upper left) and denoised cryo-EM images (right). Thon ring simulation for CTF determination is shown in lower left. (**E**) Spectra signal-to-noise ratio (SSNR) profile calculated from cryo-EM images before (orange) and after (blue) denoising. SSNR = FRC/(1-FRC), where FRC is calculated between sums of even and odd frames. (**F**) Scatter plot of defocus values determined from images before and after denoising. Defocus values were estimated using gCTF (Zhang, 2016) and the major and minor defocus values were averaged. The small population of off-diagonal images (24 of 843) appear heavily contaminated with crystalline ice.

The Fourier power spectrum (Figure 2D) and spectral SNR (SSNR) (Figure 2E) calculated from the image before and after denoising show that SNR is boosted at the low-frequency Fourier components without reduction at high frequency. This behavior is different from a linear Fourier filter (such as typically used low-pass filter), which boost low-frequency amplitude by suppressing high frequency amplitude but without any improvement in SSNR at any frequency. Thon rings associated with the CTF are present and correctly located in the denoised images, and defocus values estimated from denoised images are mostly very close to those estimated from the original noisy image (Figure 2F). A few images that have very different estimated defocus values after denoising (upper left quadrant of Figure 2F) are heavily contaminated with crystalline ice.

These examples demonstrate that CNNs trained by the *noise2noise* scheme are effective at denoising and contrast enhancement on a wide range of real single-particle cryo-EM specimens. Visualization of particles is significantly enhanced, facilitating more efficient manual image evaluation and particle picking. While we preprocess our images differently, use a novel CNN architecture, and train a single CNN model per dataset, our results are broadly consistent with other efforts of using similar denoising approaches (Bepler et al., 2019a; Bepler et al., 2019b; Tegunov and Cramer, 2019) and illustrate the robustness of the noise2noise algorithm.

### Quantitative estimation of signal enhancement and noise suppression

A major question concerning the denoising procedure we described here is whether the signal is faithfully retained at all spatial frequencies. A related question is whether the denoising procedure can facilitate any other steps in the single-particle cryo-EM pipeline beyond visual evaluation of images and particle picking.

To answer these questions, we extended the conventional cross-correlation-based estimators for SNR and spectral SNR (SSNR) to handle images modified by a deterministic, arbitrary operation such as denoising (Supplemental note 3, and related references (Baxter et al., 2009; Bershad and Rockmore, 1974; Frank and Alali, 1975)). Our approach estimates the magnitude of any ‘false signal’ or ‘bias’ added to each image during denoising. This is possible because pairs of denoised images of the same object will share signal and bias in common, while a denoised image and its paired noisy image will only share the signal. By estimating the signal variance, bias variance, and noise variance, we can compute quantities such as SNR and other similar quantities that take bias into account. Importantly, we can compute these quantities as a function of spatial frequency.

We estimated the SNR for noisy and denoised images of the entire T20S dataset (Figure 3A). While the mean SNR of the noisy images is 0.14, the mean SNR of the denoised images is 8.3. This enhancement, however, could have resulted from some noise being transformed into bias by the denoiser, inducing spurious correlations between denoised images. We define another quantity, the signal-to-noise-and-bias ratio (SNBR), which is the ratio of the signal variance and the sum of the noise and bias variances (Supplemental note 2). Intuitively, the SNBR represents the relative power of true signal compared with the power of all other components in the image. For the T20S dataset, the mean SNBR is 1.4, which is still a significant improvement over the original SNR of the noisy images.

**Figure 3.**
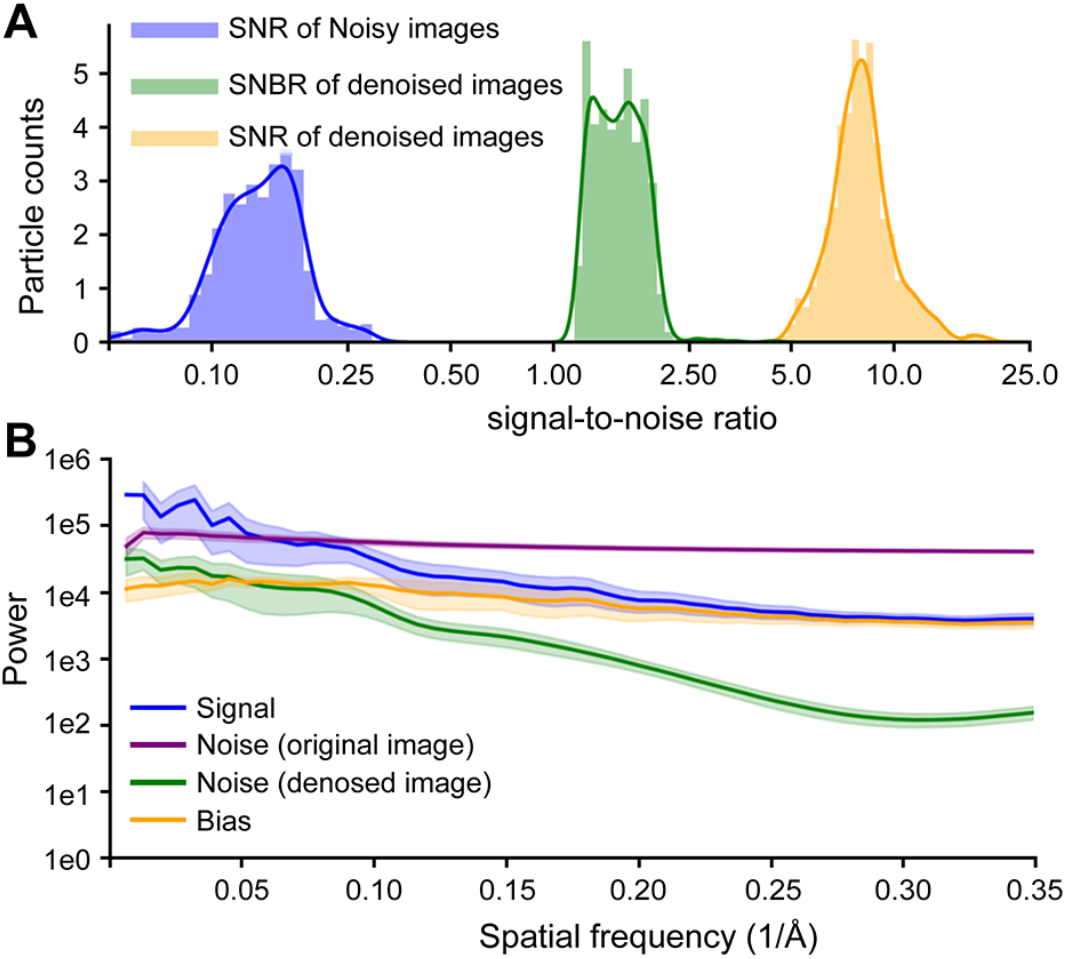
Quantitative analysis of signal, noise, and bias in denoised images. **(A)** Histograms of SNRs before (blue) and after (orange) denoising and SNBRs after denoising (green) for images from the T20S proteasome dataset. Smooth lines represent kernel density estimates of the distribution. The x-axis is on a logarithmic scale. **(B)** Spatial frequency-dependent variance (power) of the signal (blue), bias (orange), and noise. Noise power is calculated before (purple) and after (green) denoising. The y-axis is on a logarithmic scale. Curves represent the mean of the quantities for all images in the T20S proteasome dataset. Shaded regions show one standard deviation above and below each mean curve. All quantities were calculated as described in Supplemental notes 2 and 3.

We estimated the frequency-dependent variance (power) of the signal, bias, and noise before and after denoising (Figure 3B). When we plot the average of these quantities over all micrographs in the T20S dataset, we find that the noise in the original image dominates the signal in all but the lowest spatial frequencies. After denoising, the noise is much smaller than the signal at all spatial frequencies. Additionally, the signal is significantly larger than the bias at low spatial frequencies (>0.1Å^−1^) but has similar power at higher spatial frequencies.

Taken together, these results indicate that denoising increases the strength of the true signal in the denoised images by a sizable factor. However, denoising also transforms a portion of the uncorrelated noise into statistically correlated bias, which is undesirable. The nature of this bias is not clear. One possible interpretation is that very high spatial frequency patterns of signal may be impossible to disambiguate in images with very low SNR. For example, the precise arrangement of side chains in an image of a folded protein will be encoded by very high-spatial frequency Fourier components. When heavily corrupted with noise, an image of such a signal may have several plausible interpretations. Because the CNN could be wrong about any particular arrangement of side chains, the best guess for minimizing the mean-squared error is a pixel-wise average over all possible interpretations. This average would appear blurred and match no single arrangement exactly but would not be far off (in the mean-squared error sense) from any single plausible arrangement.

If this is the case, low spatial frequencies should be relatively less biased than high spatial frequencies in denoised images, because the higher SNR at low spatial frequencies should make it easier for the CNN to identify the signal unambiguously. This is what we observe in Figure 3B and is consistent with the denoising CNN enhancing the true signal at low spatial frequencies while mostly transforming the noise into bias at high spatial frequencies.

### 3D reconstructions with denoised particles

There are two questions concerning the bias being introduced into the denoised images. First, is the bias correlated between images of different particles? Assuming the correct angular orientation of each individual particle image is known, correlated bias will generate artificial structural features in the 3D reconstruction, but uncorrelated bias will be averaged out without generating artifacts. Second, would such bias interfere with the image alignment procedure and prevent structure determination directly from denoised images? We explore these two questions by performing 3D reconstructions and refinements on denoised images of the 20S proteasome.

From the T20S proteasome dataset mentioned above, we performed standard single particle cryo-EM structure determination on the original noisy micrographs, from particle picking to iterative refinement and 3D reconstruction using an initial model that is calculated from the atomic model of T20S proteasome and low-pass filtered to 60Å. The final reconstruction was refined to 3.3Å resolution from 302,290 particles using the program cryoSPARC without applying symmetry (Figure 4A, Supplementary Figure 2A blue curve). Even without sharpening, densities for side chains are clearly visible. This reconstruction as well as the orientational parameters of each particle in the final dataset are then treated as references to quantitatively evaluate the behaviors of denoised particles in both 3D reconstruction and structure refinement.

**Figure 4:**
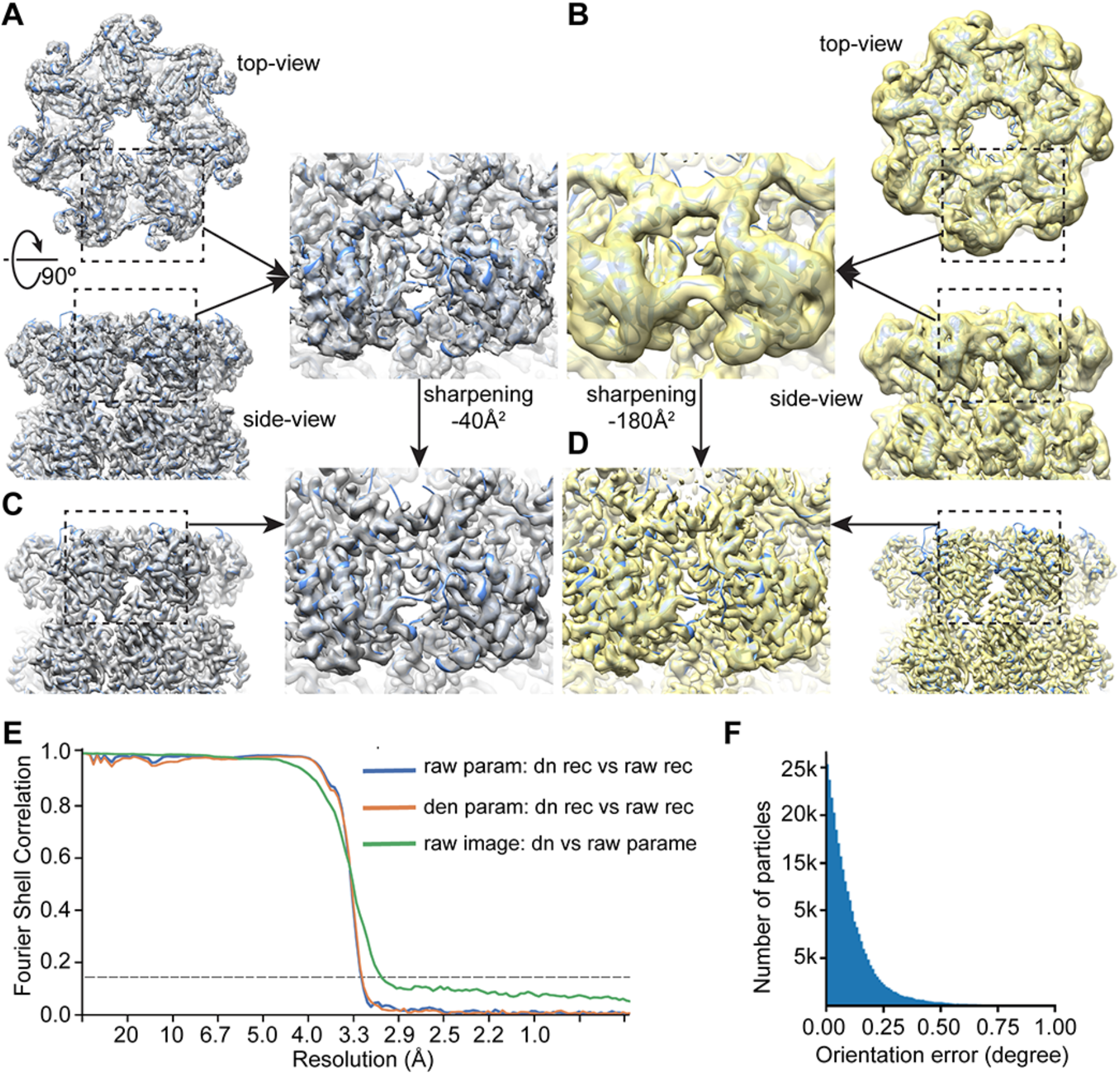
3D reconstructions of denoised 20S proteasome images. (**A**) Reconstruction without sharpening of T20S proteasome of original particle images in top (upper) and side (bottom) views. Iterative structure determination and refinement were performed using cryoSPARC(Punjani et al., 2017) with D7 symmetry. For consistency with the other panels, final reconstruction was calculated by transferring all parameters into RELION and using the program relion_reconstruct without symmetry(Zivanov et al., 2018). (**B**) Reconstruction of denoised particle images in top (upper) and side (bottom) views, with the same orientation parameters used in (A). (**C**) Reconstruction of original particles after sharpening by −40Å^2^. (**D**) Reconstruction of denoised particles after sharpening by −180Å^2^. (**E**) FSC curves between reconstructions of original and denoised particles, using orientational parameters determined from original particles (blue); between reconstructions of original and denoised particle using parameters determined from denoised particles (orange), and between reconstructions of original particles using parameters determined from either original or denoised particle (green). (**F**) Histogram of errors in the orientation parameters estimated during the refinement of denoised particles.

We extracted the same particles from the denoised micrographs and calculated a 3D reconstruction with the program relion_reconstruct by using the orientation and CTF parameters determined from the original noisy images (Figure 4B). The reconstruction of denoised particles has the correct overall shape and some detailed structural features, but appears significantly blurred compared to the 3D reconstruction of the original particles (Figure 4B), although the resolution estimation from FSC extends to 3.4Å (Supplementary Figure 2A red curve). The resolution is not uniform, as some helices are well-resolved with visible helical grooves while others are completely unresolved (Figure 4B). However, after sharpening by a negative B-factor, −40Å^2^ for the raw particle reconstruction and −180Å^2^ for the denoised particle reconstruction, all high-resolution features including side chain densities are very similar in both reconstructions (Figure 4C and D). A Fourier Shell Correlation (FSC) curve calculated from the two reconstructions is close to 1 before 4Å and falls off at 3.3Å (Figure 4E, blue curve). Because the denoised particles are band-limit to 1/3Å^−1^, this suggests that the denoised images contain high-resolution information until nearly the point where it was explicitly truncated. Importantly, it also suggests that the bias introduced by the denoising procedure is sufficiently random and can be removed by averaging large numbers of particle images. The large B-factor needed to sharpen the reconstruction of denoised particles may be caused by the significantly enhanced low-frequency SNR of denoised particle images.

We further used the same stack of denoised particles for a standard iterative procedure of particle alignment and 3D reconstruction, using the same initial reference model and RELION 3D refinement (cryoSPARC does not support refinement of phase-flipped particle images). The resolution of the final reconstruction estimated by the gold standard FSC is 3.4Å (Supplementary Figure 2B orange curve). Similarly, the reconstruction without sharpening shows strong low-resolution features, but B-factor sharpening reveals correct high-resolution features (Supplementary Figure 2C and D). The angular differences in the orientations determined from the original noisy and denoised particles are small (Figure 4F). A 3D reconstruction calculated from original noisy particles but using orientation parameters determined from the denoised particles has slightly better resolution (3.3Å, Supplementary Figure 2B green curve), and is highly correlated with the 3D reconstruction determined from the original particle images (Figure 4F, green curve). We speculate that the small angular errors are caused either by bias introduced into the denoised images or by overweighting of low-frequency information during alignment of the denoised images.

### Conclusions and discussion

The denoising CNNs we introduced here significantly enhance the contrast of noisy cryo-EM images. The most immediate applications should be visual evaluation of specimens before large-scale data collection and particle picking for small or irregularly shaped macromolecules.

Beyond these applications, quantitative evaluation of frequency-dependent signal, noise, and bias show that the true signal in the denoised images is enhanced significantly at the low and intermediate spatial frequencies required for particle alignment and maintained at high frequency.

Iterative refinement of denoised particles leads to reconstructions with nearly correct structural features at high-resolution, demonstrating the potential of using denoised particles directly for single particle cryo-EM structure determinations. Small errors in the orientation parameters determined from denoised images could be corrected by substituting and further refining the original noisy images. This reversibility is advantageous for cryo-EM image processing, unlike phase plates which improve contrast by irreversibly modulating the image.

Although we have not demonstrated it here, CNN denoising also has a potential to facilitate better identification of small classes of particles that correspond to weakly-populated intermediate states of macromolecular machines. This will be especially true if the intermediate states differ by low or intermediate resolution structural features, such as the relative positioning of a protein domain. Considering that the bias is more pronounced at high frequency, it may be desirable to merge denoised and original images in Fourier space, by combining low-frequency signal from denoised image with the high-frequency signal from the original image. We envision such merged images could be used for the entire single particle cryo-EM image processing pipeline. In the Supplementary note, we discuss a method of merging the original noisy and denoised particle images. The major impediment to directly using denoised images or merged images for alignment and classification appears to be that the current procedures implemented in cryo-EM software are not tuned to handle denoised images. This is likely caused by strong deviation of the frequency-dependent amplitude profile of denoised images compared to standard cryo-EM images.

It may also be useful to consider other uses for denoised images. With higher SNR single particles, the particle is clearly delineated from the background. This would make per-particle real-space masking possible for small particles with irregular shapes, eliminating most of the noise surrounding the particle and presumably enhancing alignment. Similarly, it could make previously proposed pseudo-atom approaches for estimating initial models and measuring macromolecular flexibility more tractable (Joubert and Habeck, 2015).

## Acknowledgment

We thank Drs. Joshua Batson, David Dynerman and David Agard for thoughtful discussions. This work is supported in part by NIH grants R01GM098672, R01HL134183, P50AI150476, P01GM111126, S10OD020054, and S10OD021741 to Y.C. Y.C. is an Investigator of Howard Hughes Medical Institute.

## Supplemental note 1: Mathematical description of the *noise2noise* training scheme

First, we assume an image (M) to be a vector of pixels composed of a signal vector (S) and an additive noise vector (N). Though we do not explicitly denote it, the noise component may or may not depend statistically on the signal and therefore can include shot noise (Poisson noise).

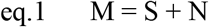

A parameterized denoiser is a function with parameters θ that takes the noisy image, M, and outputs the signal, S:

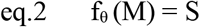

The conventional strategy for training a deep CNN to be an image denoiser is to search for the set of parameters, θ, that minimizes the expected value of a loss function (the expected risk) over all the clean/noisy image pairs in the training data:

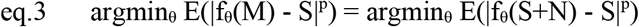

where E(X) denotes expectation over the data distribution and |X|^p^ denotes the L_p_ norm over all pixels in the resulting difference image. In this and other related works, p=2 and corresponds to the mean-squared error over pairs of corresponding pixels, though L_1_ (median-seeking) and approximate L_0_ (mode-seeking) norms were also demonstrated in Lehtinen et al. These parameters are determined using some variant of stochastic gradient descent over many small batches of the training data. Because f_θ_ (and any CNN in general) is a differentiable function, the gradient of each parameter with respect to the loss can be estimated directly using the chain rule (the back-propagation algorithm).

The key insight of Lehtinen et al. is that one can learn the same parameters without clean data. Instead of M and S, assume we have two images with the same signal but uncorrelated noise:

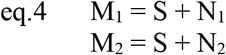

We make two very general assumptions about the noise:

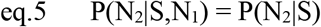

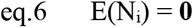

where P(N|S) is the conditional probability distribution of the noise given the signal, E(N_i_) is the expectation over the distribution of noise, and **0** is a vector of zeros with the same shape as the images M_1_ and M_2_.. Intuitively, these conditions imply that the noise present in an image pair is statistically independent from each other given the signal and that the noise is zero-mean and ‘averages out’ if many noisy images are summed together. Then, we train the CNN with stochastic gradient descent or one of its variants to minimize the following objective:

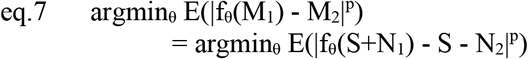

In this procedure, while we are training f_θ_ to convert one noisy image, M_1_, into the second noisy image M_2_, because of (eq.5), M_1_ provides f_θ_ with *no information* about the specific instantiation of noise N_2_ that it is tasked to predict: N_2_ is an independent random draw from the noise distribution P(N|S). To minimize the expected risk, the best guess f_θ_ can make for N_2_ is the average of all the possible instantiations of noise it might see, which by (eq.6) is zero noise. This implies that the objective in (eq.7) is minimized by the same set of parameters **θ** that minimizes (eq.3), and so (eq.7) trains f_θ_ to be a denoiser.

The *noise2noise* strategy is more intuitive when considered geometrically, when an image is represented by a vector of pixel intensities, locating a single point in a high-dimensional vector space. A signal, S, occupies a point in the space. Noise is a vector N_i_ that displaces an image away from signal S to a noisy image S+N_i_. The conventional denoising CNN strategy trains a mapping between a given noisy point S+N_i_ and S. The *noise2noise* strategy attempts to train a mapping between a given noisy point S+N_1_ and a second noisy point S+N_2_, but provides no information about N_2_. Because N_2_ could be any of the possible noisy points and we penalize f_θ_ if its output is distant from S+N_2_, the least risky guess is the point that is closest to all of the noisy points. This point is simply the mean of all noisy images S+N_i_, which is also the noiseless signal, S.

## Supplemental note 2: Technical description of the training and denoising implementation

Code implementing these methods is available at https://github.com/eugenepalovcak/restore.

All code was written in python. Neural networks were implemented and trained using the libraries keras and Tensorflow (Abadi et al., 2016).

### I/O and image pre-processing

Our program, *restore*, takes as input a RELION-style STAR file containing the file path and CTF parameters for a set of training micrographs. Each image in this STAR file is assumed to be the sum of all frames summed together after motion correction and dose-weighting with MotionCor2 (Zheng et al., 2017) and CTF estimation with CTFFIND (Mindell and Grigorieff, 2003), Gctf (Zhang, 2016), or some other CTF-estimation program. Each training image is expected to have a training pair of even/odd images. These can be generated with recent versions of MotionCor2 using the ‘-SplitSum 1’ option. In this case, the full sum image has the suffix ‘DW’ while the even/odd half sum images have the suffixes ‘EVN’ and ‘ODD’.

*restore* begins by preprocessing the training data. For each even and odd half-sum image, we load the MRC file into an array. Then, we Fourier crop the micrograph such that the pixel size is ½ the value of the ‘--max_resolution parameter’, effectively setting the nyquist frequency ‘--max_resolution’. A band-pass filter is applied in Fourier space to remove low frequency information (lower than 1/200 angstroms, corresponding to gradients of image contrast due to changes in ice thickness) and high-frequency information beyond the Nyquist frequency.

Next, we correct for the CTF by phase-flipping. Explicitly, we calculate the amplitude modulation implied by the CTF for each Fourier component, compute the sign of each amplitude modulation (+1 or −1), and multiply the Fourier transform of the Fourier-cropped image by this array. We inverse Fourier transform this image. It is worth noting that we also tried denoising without phase-flipping and found similar results as with phase-flipping. However, we did not extensive test this option.

Then, we break each even and odd half-sum image up into 192×192-pixel patches. We normalize by subtracting the mean and dividing by the standard deviation. We store the patches in a large hierarchical data format (HDF) file. This allows fast access to the preprocessed data during training without requiring the entire dataset to be loaded into memory.

### CNN architecture

We specified the architecture of the CNN using the keras library. Our CNN has a U-net architecture with specialized CNN block structures. The encoder branch consists of an initial 2D convolutional layer followed by three larger blocks of convolutional layers that progressively down-sample the input. In the decoder branch, three blocks of convolutional layers then progressively upsample the feature maps and receive ‘skip connections’ from the encoder.

The blocks of convolutional layers are modeled after the wide-activation super resolution (WDSR) networks described by Yu et al. The basic building block of these networks is the wide-activation convolutional layer. Wide activation convolutional layers take linear combinations of feature maps to increase their depth before applying a non-linearity and convolution operation. These expanded linear combinations allow more information to pass through the layers without getting decimated by the ReLU non-linearity (the so-called ‘vanishing gradient’ problem). For the same reason, each layer also uses residual connections, adding the processed information to the input.

Wide-activation convolutions are implemented by performing a 1×1 convolution to increase the depth of the input feature maps from 32 to 128, applying a ReLu non-linearity, and performing another 1×1 convolution to reduce the feature maps from 128 to 25. Finally, a 2D convolution is applied with 32 feature maps and 3×3 kernel size. A down-sampling wide-activation block consists of two branches. One performs a simple max-pooling operation. The other contains 4 wide activation convolutional layers followed by a down-sampling 2D convolution with stride 2. The feature maps of each branch are summed. An up-sampling wide-activation block also consists of two branches. The first consists of a single upsampling layer, which expands the input feature map depth from 32 to 128, applies a wide-activation convolution, and applies the depth-to-space upsampling operation. The other branch concatenates the input feature maps with those from encoder’s skip-connection. These feature maps are then passed through 4 wide activation layers and an upsampling layer. The upsampled feature maps from both branches are then added together.

The final layer of the decoder branch is a 1×1 convolution operation that predicts the pixel values of the output image. In total, the full CNN model consists of 96 convolutional layers with 2.3 million trainable parameters.

### Training and applying the CNN

To train such a large CNN on consumer GPUs, we used several tricks to reduce memory consumption. First, instead of using batch normalization to stabilize training, we used weight normalization. This enabled us to use smaller training batches (10 patches per batch) without training becoming slow or erratic. Second, we used gradient checkpointing to reduce GPU memory usage at the expense of longer computation time. We trained the network using the Adam stochastic optimizer for 100 epochs of 500 batches each. We initialize the learning rate at 1E-4 and decrement it by ¼ every 25 epochs.

Once trained, image paths and CTF parameters are read in from a STAR file. Each micrograph is then denoised, beginning with the same preprocessing pipeline used to generate training data. Instead of dividing the image into patches, the whole downsampled, phase-flipped image is passed through the trained CNN. Then, the image is padded with zeros in Fourier space and and a soft low-pass filter is applied at the Nyquist frequency of the down-sampled image. When inverse Fourier transformed, the final denoised image has the same pixel size and dimensions as the noisy input image and has the CTF corrected by phase-flipping.

### Merging denoised and noisy images

Merging with noisy image and the denoised image is performed by high-pass filtering the noisy image and low-pass filtering the denoised image with complementary soft-edged Fourier filters. Instead of Fourier padding into an array of zeros, the denoised image is effectively Fourier padded with the high-frequency information from the original noisy image. If not properly normalized, sharp changes in amplitude can occur in the band of spatial frequencies where the highpass and lowpass filters overlap (the merge band). To avoid this, the amplitudes of the denoised image are scaled by a normalization factor that keeps the standard deviation of amplitudes in the merge band the same after merging the denoised image into the noisy one. The frequency of the lowpass filter and the spectral width of the soft edge (both in angstroms) are parameters the user can specify.

MRC I/O operations use the library *mrcfile*. STAR file/CTF operations use our library pyem. All other image preprocessing is done using numpy, particularly the real FFT/IFFT routines. All CNN operations were performed with the library keras using the Tensorflow backend.

### Measuring the angular errors between pairs of orientation parameters

Ignoring the in-plan x-y positions of particles, we measured the angular distance between orientations of each denoised and raw particles determined by separate refinements by using a quaternion representation for the rotation (explicitly, the inverse cosine of the norm of the quaternion inner product). The D7 symmetry of the T20S proteasome was taken into consideration when calculating these angular differences. For all quaternion-based calculations, we used the *geom* module of the library *pyem* (Asarnow et al., 2019).

## Supplemental note 3: Estimating signal, noise, and bias in denoised cryo-EM images

Once we have denoised cryo-EM images with a trained CNN, we would like to evaluate the SNR of the denoised image and the magnitude of any systematic errors added by the denoiser. Additionally, we would like to estimate these quantities as a function of spatial frequency.

SNR is defined as the ratio of the signal variance and the noise variance (Bershad and Rockmore, 1974; Frank and Alali, 1975):

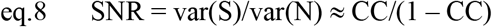

Here, var(S) and var (N) are variance of signal and noise, and CC are cross correlation between two images of identical object. Classic approach of using cross-correlation based estimators to calculate SNR require a pair of images taken from an identical object under the identical optic conditions. Historically, it was prohibitively inconvenient to obtain two cryo-EM images of an identical target, so quantitatively estimating the SNR or the spectrum SNR (SSNR) was impossible. Instead, SNR of cryo-EM image was estimated from the Fourier power spectrum, where the background noise spectrum is estimated by fitting a smooth function through the CTF zeros (Booth et al., 2004). This makes the assumption that the oscillating Thon rings in the power spectrum result exclusively from the signal, whereas noise is unaffected by the CTF. Given the power spectrum and background noise spectrum, the SSNR can be estimated as ((P(s) – N(s))/N(s) (Booth et al., 2004). While it is straightforward to calculate the SSNR from the Fourier power spectrum in this way, it is impossible to estimate or to remove the influence of the bias introduced by the denoising procedure, as bias may also be modulated by the CTF.

However, modern cryo-EM images are collected with high-speed direct electron detectors as stacks of frames. Assuming that the noise is independent in each frame (as we would expect for shot noise), it is possible to estimate the SNR and SSNR using the classic two-image cross-correlation approach (eq. 8) by calculating two sums of odd and even frames.

We assume that a denoised image (D) is the sum of the signal (S), the remaining uncorrected noise (N_d_), and some false signal (B) that results from systematic error in the denoiser (the bias of the estimator). We distinguish N_d_ from the noise in the original image, which we refer to as N_n_.

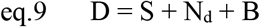

Thus, we need to estimate var(S), var(N_d_), and var(N_n_). We also would like to estimate var(B) for the denoised images. We recall several elementary identities of the variance and covariance:

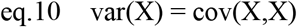

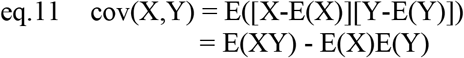

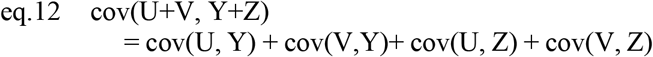

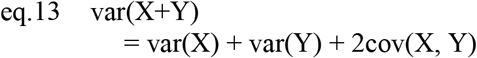

eq.13 is implied by eq.10 and. eq12. If X and Y are statistically independent, E(XY)=E(X)E(Y) and eq.11 implies that cov(X, Y)=0.

Computationally, the covariance of two images can be estimated with:

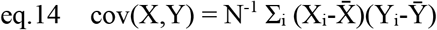

Here, N is the number of pixels in images X and Y, X_i_ is the intensity of the i^th^ pixel of image X, and 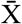 is the mean pixel intensity.

We denote the noisy even and odd image sums as M_1_ and M_2_ and their denoised versions as D_1_ and D_2_. From the previous equations, it is straightforward to show with arithmetic that:

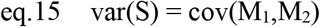

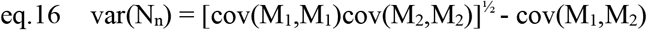

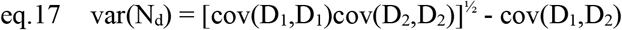

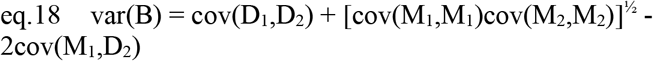

In eq.16, the first term on the right-hand side is var(S+N), estimated as the geometric mean of var(M_1_) and var(M_2_). Similarly, the first term on the right-hand side of eq.16 is var(S+N+B) and is estimated as the geometric mean of var(D_1_) and var(D_2_). From these equations, we can estimate the SNR of the images before and after denoising:

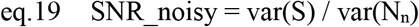

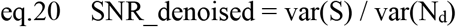

Intuitively, eq.19 is identical to the expression for the SNR of noisy image pairs used by the Frank and Alali (Frank and Alali, 1975) and originally suggested by Bershad and Rockmore (Bershad and Rockmore, 1974):

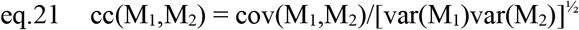

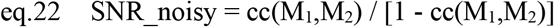

We can also estimate other potentially interesting quantities such as the ratio of the signal variance and the bias variance (signal-to-bias ratio, SBR).

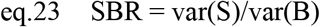

Finally, we can estimate each of these quantities as a function of spatial frequency. To estimate the covariance of X and Y at some spatial frequency k, we calculate:

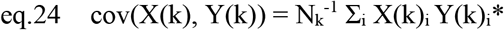

where X(k) is the ring of Fourier components in the Fourier transform of X with spatial frequency k, N^k^ is the number of Fourier components in ring k, Y(k)* denotes the complex conjugate of Y(k), and the sum runs over each corresponding pair of Fourier components, i. This approach treats each ring in the Fourier transforms of X and Y as an independent random variable.

---

**Supplementary Figure 1:**
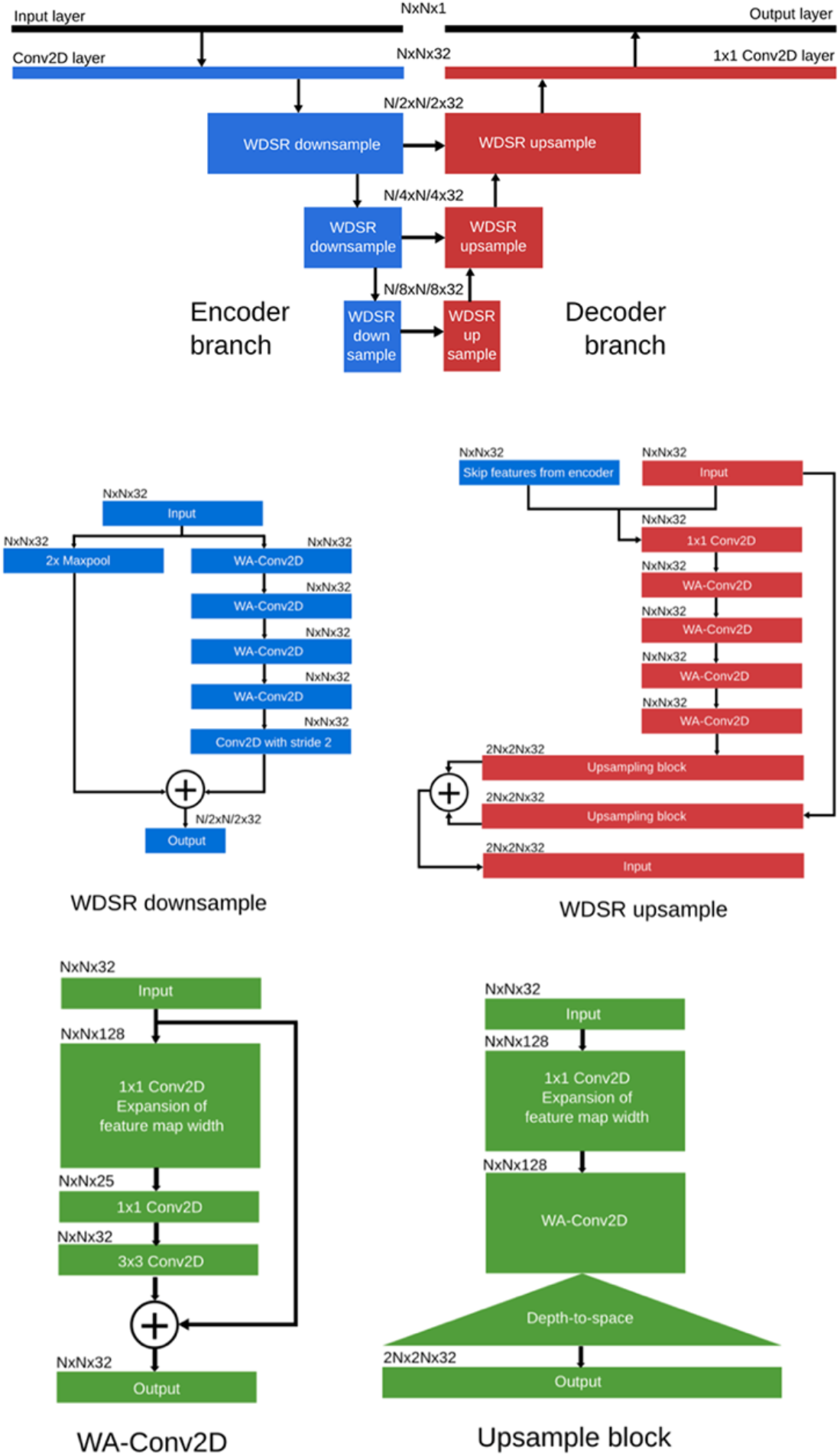
Architecture of the convolutional neural network. (A) The network has an encoder-decoder structure, similar to the U-net architectures commonly used in biomedical image segmentation. Each convolutional downsampling or upsampling operation in the CNN is performed by a ‘wide-activation super-resolution’ (WDSR) sub-network. (B) The WDSR down-sample subnetwork (left network) consists of 4 wide-activation convolutional layers (WA-Conv2D, described in C) followed by a conventional convolution with stride 2. The resultant downsampled feature maps are summed with the input feature maps, which are downsampled by 2x max-pooling, as in a residual neural network. The WDSR up-sample network (right) first concatenates the input feature maps and the feature maps from the skip connections and reduces them with a 1×1 convolution operation (N×N×64 → N×N×32). After 4 WA-Conv2D operations, the feature maps are 2x upsampled with an upsampling block (described in C). Similar to the downsampling network, the input feature maps are upsampled and summed with the upsampled output feature maps. (C) The WA-Conv2D operation (left) first uses a 1×1 convolution operation to ‘expand’ the input feature maps into a higher dimensional space (N×N×32-->N×N×128). Intuitively, each feature map of the higher dimensional space is a distinct linear combination of the input feature maps. A ReLU non-linearity is applied to the expanded feature maps, followed by another 1×1 convolution operation to reduce the feature map depth. A conventional convolution operation is then applied with a 3×3 convolutional filter. The upsampling block (right) consists of a WA-Conv2D operation followed by a depth-to-space upsampling operation.

**Supplementary Figure S2:**
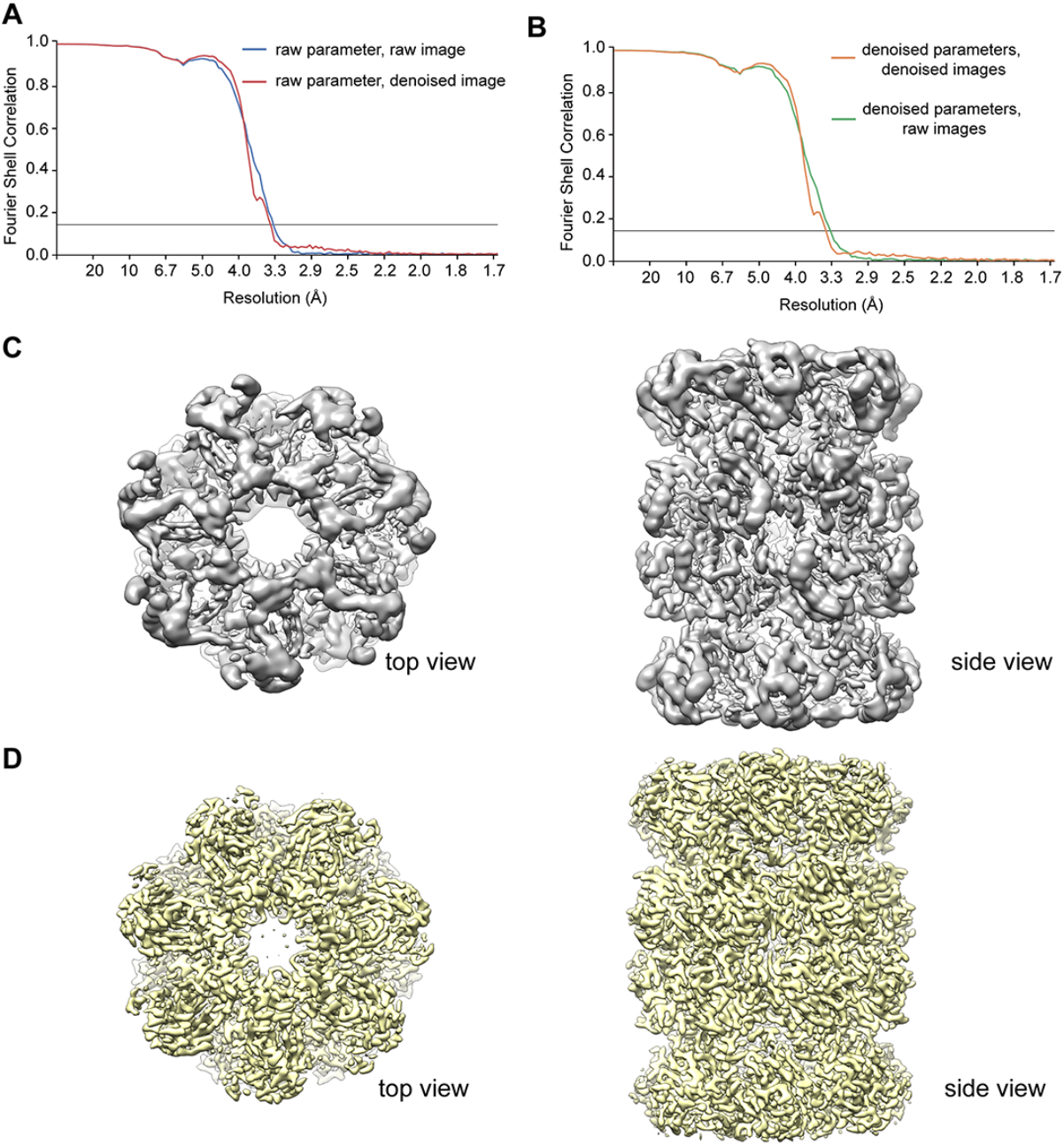
(**A**) Gold standard FSC curve between two half maps of T20S proteasome determined from original noisy particle images (blue) and FSC curve between two half maps calculated using the same parameters but denoised particle images (red). (**B**) Gold standard FSC curve between two half maps of T20S proteasome with orientation parameters determined from denoised particle images (orange) and FSC curve between two half maps calculated using the same parameters but original noisy particle images (green). (**C**) Side and top view of 3D reconstruction determined noisy images, before B-factor sharpening. (**D**) same views of 3D reconstruction sharpened by a B-factor of −180Å^2^.

